# Protozoan predation drives adaptive divergence in *Pseudomonas fluorescens* SBW25; ecology meets experimental evolution

**DOI:** 10.1101/2021.07.12.452127

**Authors:** Farhad S. Golzar, Gayle C. Ferguson, Heather Lyn Hendrickson

## Abstract

Protozoan predators can affect the structure of bacterial communities, but investigations of how predation might influence bacterial evolution and antagonistic behaviours are scarce. Here, we performed a 20-day predator-prey evolution experiment on solid media to investigate the effect of continuous protozoan predation on bacterial traits using *Pseudomonas fluorescens* SBW25 as prey and *Naegleria gruberi* as an amoeboid predator. We observed the divergence of colony morphotypes coincident with an increase in bacterial grazing resistance and relative prey fitness in selected bacterial isolates. When subjected to these resistant prey, *N. gruberi* show reduced activity (increased encystment) and limited replication. An investigation of the mutations responsible for predation resistance reveals mutations in *wspF* and amrZ genes, affecting biofilm formation and motility. The bacterial mutants in the *wspF* gene successfully colonise the air-liquid interface and produce robust cellulose biofilms that prevent predation. The mutation in the *amrZ* mutant withstands predation but this variant produces low levels of cellulose and limited swarming motility. Our findings suggest that protozoan predation can profoundly influence the course of genetic and phenotypic evolution in a short period.

## Introduction

The microbial world is replete with examples of competition and predation. These microbial melees impact the traits that bacteria evolve. Ultimately, the bacterial phenotypes that are successful in these contests affect agriculture, the environment and human health [1–3]. The predators of the microbial world include the free-living amoebae (FLA), single-celled eukaryotic protozoan predators that consume bacteria by phagocytosis and are ubiquitous in diverse environments [4]. Protozoan predation asserts a strong selective pressure on bacteria, reportedly eliminating 60% of bacteria in soils [5–8]. Despite their effects on decreasing bacterial abundance, FLAs are often neglected in considerations of microbial ecology [9,10]. They are, however, powerful agents involved in decomposition, recycling nutrients, and energy in ecosystems [1–3, 11–13].

FLA predation is a potent selective force driving bacterial evolution [14]. Failure to resist predation results in whole-cell ingestion and digestion of bacterial prey [15]. Amoebae are however inefficient and careless predators [16,17]. Bacteria can escape protozoan predation by adopting anti-predatory behaviours such as cell elongation, secondary metabolite or toxin secretion, and mucoid phenotypes [14,18–24]. Investigating how predation drives bacterial diversity is fundamental to understanding how microorganisms evolve in the environment.

FLAs have been proposed to be a major driving force in the evolution of traits such as biofilm formation, intracellular growth and encapsulation in bacteria [25–28]. These complex sets of anti-predatory strategies benefit bacteria in the face of protozoan predation and help to maintain stable communities [18]. Biofilm formation is a common adaptation in prokaryotes contributing to survival in many different environments [29,30]. Growth in a multi-cell extracellular matrix affords protection from predatory phagocytosis whilst allowing bacteria to colonise a range of otherwise hostile environments [31,32], attaching to surfaces, and invading new niches [32–34]. In the presence of the amoeboid predator *Acanthamoeba castellanii, Pseudomonas aeruginosa* cells have been demonstrated to form protective biofilms, making the bacteria non-edible [28,35,36]. Biofilm formation can also be beneficial through the exclusion of competing organisms [37], and can increase resistance to harsh environmental conditions like antibacterial agents [38], metal toxicity, acid, and salinity [39,40].

In *P. fluorescens* SBW25 biofilm formation is a well-studied adaptation to oxygen limitation induced when growing at the air-liquid (AL) interface in standing liquid cultures [41,42]. In this system, mutations that induce the constitutive expression of extracellular polysaccharides are well documented (wsp mutations) [42,43]. Wrinkly Spreader (WS) phenotypes are biofilm forming mutants that arise from the ancestral Smooth (WT) colony morphotype, allowing access to the oxygen rich surface of the liquid culture. Fuzzy Spreaders (*fuzY* mutations) are early colonisers of the surface that ultimately occupy the bottom of the microcosm [44–48].

Amoeboid predation has also been hypothesised to select for bacterial virulence as early as 1988 [49]. This hypothesis, sometimes called the “coincidental evolution hypothesis” suggests that adaptations to one environment, like protozoan predation, could result in virulence traits expressed in a similar environment, like a metazoan host. While tantalizing correspondences have been observed between virulence and predation resistance, a direct test of this idea has not been attempted [14]. These ideas have recently experienced renewed interest and experimental attention [29,49,50]. Many pathogens have traits that deter predation. For example, *Klebsiella pneumoniae* cells are capable of producing lipopolysaccharide (LPS) and outer membrane proteases that limit amoeboid predation by *Dictyostelium discoideum* [51].

Whilst there is abundant literature describing a range of bacterial traits that may have evolved as anti-predatory devices, there is little or no experimental work that observes the traits that bacteria acquire in response to the pressure to survive FLA predation. In order to investigate the effect of continuous FLA predation on bacteria we used an experimental evolution approach in which populations of *P. fluorescens* SBW25 were continuously preyed upon by a non-pathogenic amoeba, *N. gruberi* NEG-M, on solid surfaces. We hypothesised that traits associated with virulence would arise in response to our protozoan predation regime.

## Materials and Methods

### Strains and Media

A strain of *P. fluorescens* SBW25 (NC_012660.1) that constitutively expresses GFP was used in order to easily confirm the genetic background of the strain and detect contamination [52]. The strain was regularly grown in 5 mL Lysogeny Broth (LB) [53] from −80°C frozen stock and incubated at 28°C in a shaker (180 rpm). Protozoan Strain *N. gruberi* NEG-M was obtained from the American Tissue Culture Collection (ATCC^®^ 30224™) and was originally derived from strain NEG (isolated from soil in 1967 in Berkeley, California). *N. gruberi* cells were propagated on solid PM media (4 g Difco Bacto Peptone, 2 g dextrose, 1.5 g K_2_HPO_4_, 1 g KH_2_PO_4_ and 20 g agar (2%) per litre H_2_O) supplemented with 2% heat-inactivated Fetal Bovine Serum (FBS) (MEDIRAY, MG-FBS0820) and incubated in parafilm sealed plates at 28°C [54].

For the purpose of plaque assays (Plaque Forming Unit) (Fig. 2A) in which only WT was present, PM media was supplemented with 33 mg of cholesterol (Sigma-Aldrich, C8867) per liter of PM agar in parafilm sealed plates and incubated at 28°C [55]. This was sufficient to allow for growth of *N. gruberi* on the ancestral WT strain in order to measure PFU. However, in all other experiments, FBS was supplemented, as described above. *N. gruberi* stock was regularly stored in a 2 mM Tris HCL buffer (121.1 g Tris base in 800 mL H_2_O, 60 mL HCl to get pH 7.6 (autoclaved or filter sterilised) at room temperature [56]. Amoebae were cultivated on WT bacterial prey before experimental evolution on medium-sized Petri plates (90 x 15mm).

### Prey-Predator Experimental Evolution

Experimental evolution was performed on solid PM agar plates, supplemented with 2% FBS to support long term *N. gruberi* growth. Three replicates of prey-only treatment were used as a control by spreading a lawn of 1×10^8^ *P. fluorescens* SBW25 cells in 100 μl on agar plates. Co-evolution of prey and predator were started in the same way with the subsequent addition of 1×10^3^ protozoan cells in 2 μl of Tris HCl buffer on one side of the agar plate on a sterilized filter disk. Plates were wrapped in parafilm to prevent drying of the medium and incubated at 28°C to allow *N. gruberi* to consume the bacterial lawn across the plate.

Predation plates were developed for 5-day cycles, after which the lawn of the surviving prey bacteria and protozoan predators or prey-only controls were each washed with 3 mL of Tris buffer. The wash from the co-evolution plates was subjected to centrifugation at 400 g for 3 minutes in order to separate the amoebae (primarily in their encysted state) from bacterial cells. The supernatant was removed and 100 μl of Tris HCL was used to resuspend the amoebae pellet. Approximately 3% (100 μl) of the supernatant, and 2% (2 μl) of the resuspended pellet containing protozoan cells was transferred to fresh solid FBS supplemented PM plates for the next cycle. This cycle was performed every 5 days (4 cycles) for a total of 20 days. The remaining bacterial cells were frozen in equal parts for a final concentration of 4.25% NaCl and 35% glycerol at −80°C for further investigation. We were unable to reliably cryopreserve *N. gruberi* during the course of the experiment.

### Isolation of the Predation Adapted Bacteria

After 20 days of evolution the surviving bacterial cells were plated for colony-forming units (CFU) and novel colony morphologies were selected for single colony isolation on LBA plates containing 100 μg/mL ampicillin and preserved at −80°C as above for further study. In addition, three smooth colonies from the prey-only plates were selected for single colony isolation and similarly preserved.

### Predator Growth on Bacterial Isolates

The ability of predator-evolved bacterial isolates to affect the viability of amoebae was established by growing ancestral *N. gruberi* on evolved bacterial isolates and comparing PFU on these novel bacteria to PFU on WT *P. fluorescens* SBW25. Bacterial cultures were first grown overnight in 5 mL of LB in a shaker (180 rpm). The ancestral stock of *N. gruberi* was diluted in order to yield 5-10 plaques on a total of six replicate plates [54]. Briefly, protozoan cells were transferred into the tubes that contained overnight test cultures of ~1×10^8^ bacteria (100 μl), mixed thoroughly and spread onto PM agar plates. Plates were wrapped in parafilm and incubated at 28°C for four days. Plates were monitored daily for protozoal plaque formation and recorded (Fig. 2 A).

To evaluate the number of *N. gruberi* generations (n) on each coevolved bacterial population, bacterial test isolates of interest were mixed with protozoan cells (10 +/- 2) and incubated as described above. The initial number of *N. gruberi* cells placed on each plate was estimated by counting PFU at T=0 (ci). Protozoan cells were grown on predation-evolved bacterial isolates of interest for four days of growth at T=4 (cf). These plates were subsequently washed with 1.5 mL of Tris HCl (7.6 pH) buffer and plated for PFU as described above.

The average number of generations that amoebae achieved on the various prey mutants was then calculated as:

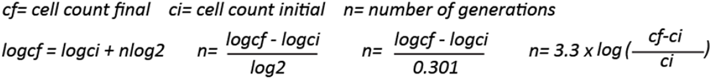

### Predator Survival Estimates

In order to determine the numbers of live or dead amoebae in either the encysted or amoeboid states after predation on the predator-evolved bacterial isolates, we performed flow cytometry. Briefly, 1×10^3^ ancestral protozoan cells were placed on one side of the PM plates and bacterial isolates were streaked with a 1 μl sterile loop from the side of the plates downwards. After 2-3 days of growth, interaction zones containing protozoan cells were gently harvested using a 1 μl loop and resuspended in 10 μl of Tris HCl buffer and stained with 2% Propidium Iodide (V/V) (Thermo Scientific, P3566). PI stains cells that have lost membrane potential. Cells were incubated at room temperature for 10-20 minutes in the dark and examined by flow cytometry (BD FACSCanto™ II) using a PE-A channel. The ratio of stained to unstained cells in an appropriate gate was used to determine the proportion of the population that was not viable.

### Line Test Assay

In order to measure the rate of predation by ancestral *N. gruberi* on the various bacterial isolates, line test assays were conducted as previously described [57]. Briefly, overnight cultures of six evolved isolates and the *P. fluorescens* SBW25 WT control were grown in LB from −80°C glycerol stocks. Approximately 1×10^3^ protozoan cells were added to a sterile filter paper placed in the middle of the PM plate supplemented with FBS. Bacterial isolates of interest were streaked as replicated paired lines on the plate with a 1 μl loop, from the centre of the plates outwards (Fig. 3A). Plates were wrapped in parafilm and incubated at 28°C for five days. Plates were photographed every 24 hours and the distance of protozoan consumption of the bacterial lines was recorded.

Predation rates were calculated as the average of the rate of line disappearance over time for each evolved type over at least six replicate lines. These were adjusted for absolute bacterial density based on relative CFU of plugs of the plates. Prey fitness values were calculated by multiplying the overall rate of predation (mm^2^/day) by the normalized cell density (cells/mm^2^), normalizing relative prey fitness (cells/day) to set the value of the least-preferred mutants (WS1, FE, & WFE) to 1 and the most preferred mutant to 0 (LWT1) [57].

### Bacterial Fitness Assay

Overnight cultures of bacterial isolates were grown in LB from −80°C glycerol stocks. The bacterial fitness assays were performed on solid PM agar plates. To estimate the growth rate of bacterial lines in the absence of protozoa, replicate cultures of 1×10^7^ bacteria (30 μl) were spread on small Petri plates (35 x10 mm). For the predation group, 1×10^7^ bacteria (30 μl) were spread in the same way and quantities of ancestral *N. gruberi* cells (1×10^3^) were subsequently added to one edge of each plate. All plates were wrapped in parafilm and incubated at 28°C. After four days, plates were washed with 1.5 mL Tris HCl buffer, serially diluted, and plated for CFU. The average number of bacterial generations was estimated by obtaining the initial and final number of bacteria placed on each plate (i.e the CFU at T=0 and T=4, respectively).

### Motility Test Assy (Bacterial Swarming)

Bacterial swarming was examined on 0.5% agar M9 plates (100 mL M9 salt, 1 mL 1M MgSo_4_, 10 mL 20% glycerol, 2.5 grams casamino acid and 50 μl 1M CaCl_2_, adjusted to 500 mL with H_2_O). Bacterial isolates were grown overnight in LB from frozen glycerol stocks. Assays were conducted by dipping the overnight bacterial culture and stabbing into the surface of agar plates using 1 μl inoculation Loops (n=10). Plates were incubated for 72 hrs at 28°C and photographed when required.

### Microscopy and Image Analysis

#### Amoebae and bacteria visualisation

cells were harvested from actively growing PM plates using a 1 μl loop placed and distributed gently on a glass microscopic slide on an agarose pad (M9 media +1% agarose). Cells were allowed to dry and were then covered with a glass cover slip. Images were acquired using PH and GFP filter channels (excitation 80 and 250 nm, respectively) with a 100x phase objective lens (Olympus BX51).

#### Cellulose staining

cell microscopy was performed using a fluorescence inverted microscope (Nikon Eclipse Ti2). Samples were grown overnight from glycerol stocks in 5 mL of KB broth [58]. From the overnight culture, 5 μl was dropped into fresh KB broth containing 200 μg mL^-1^ calcofluor white (Fluorescent Brightener 28, Sigma-Aldrich) and plates were incubated overnight at 28°C. Portions of the resulting bacterial colonies were sampled by scraping and placed directly on a glass microscope slide and covered with a glass coverslip. Images were captured using the Dapi filter channel (excitation 380 nm) using a 100x phase objective lens.

#### Capsule staining

Capsule visualisation (Sup. Fig. 2) was performed by growing the isolate of interest overnight in KB broth and mixing 10 μl of the culture with 10% India Ink (V/V) (PE316000). 2-5 μl of the mixed solution was then placed onto one side of the microscopic slide, thoroughly spread using the edge of the coverslip with an angle of 45° and allowed to dry for 5-10 minutes at room temperature. Slides were then viewed using phase contrast with a 100x phase objective lens (Olympus BX51).

#### Colony images

Colony morphology images were carried out using a Zeiss dissecting microscope (Stemi 2000-C). Phase contrast images were acquired using SwiftCam x 0.5 and 1.2 objective (MA95011). Cell and colony images were processed in ImageJ as described in figure legends.

### Microbial Mat Strength Assay

To compare the strength of the biofilms across the various mutants that were isolated, a mat strength assay was performed as previously described [44]. Bacterial isolates of interest first were inoculated from frozen stocks kept at −80°C and grown overnight in 5 mL KB broth at 28°C. 5 μl of the overnight test isolates in four replicates were diluted into fresh 5 mL KB microcosms. Evolved bacterial isolates were incubated at 28°C in a static environment for 72 hours. Bacterial isolates were then tested for the strength of biofilm after 3-day growth in KB by gently placing the maximum number of 2 mm glass beads in the centre of the biofilm until the mat collapsed in the microcosm vial.

### Whole-Genome Sequencing of the Bacterial Isolates

Representative isolates were revived from frozen −80°C stocks. DNA extraction was performed as described for whole-genomic DNA purification using the Promega™ Wizard™ Genomic DNA Kit (A1125, Promega). Genome quality and quantity were checked in 1% agarose gels stained with %0.0001 SYBR Safe and measured in a NanoDrop (ACTGene ASP-3700, Alphatech Systems), respectively. Genomes of all coevolved, prey-only controls, and ancestral *P. fluorescens* SBW25 WT were sent to MicrobesNG (www.microbesng.com) for 250 bp paired-end, next-generation Illumina sequencing. Sequenced reads were trimmed, aligned, and mapped to the *P. fluorescens* SBW25 reference genome (NC_012660.1) [59] for a minimum and maximum sequencing depth of 60 and 190, respectively, across the genome using GENEIOUS version 9.0.5 [60]. Mutations in isolates were identified as being present in 90% of reads in the alignment by GENEIOUS using the “Find Variation/SNPs” tool.

## Results

### Short Term Experimental Evolution of Bacteria under Predation by Protozoa

In order to test the degree to which amoeboid predation contributes to the evolution of bacterial virulence, we established a short term co-evolution experiment on solid media (Fig. 1A). A GFP labelled strain of otherwise WT *P. fluorescens* SBW25, a non-pathogenic plant-associated bacterium, was subjected to continuous predation by common soil-based amoeboid protozoan predator *N. gruberi* NEG-M (Fig. 1B).

**Figure 1.**
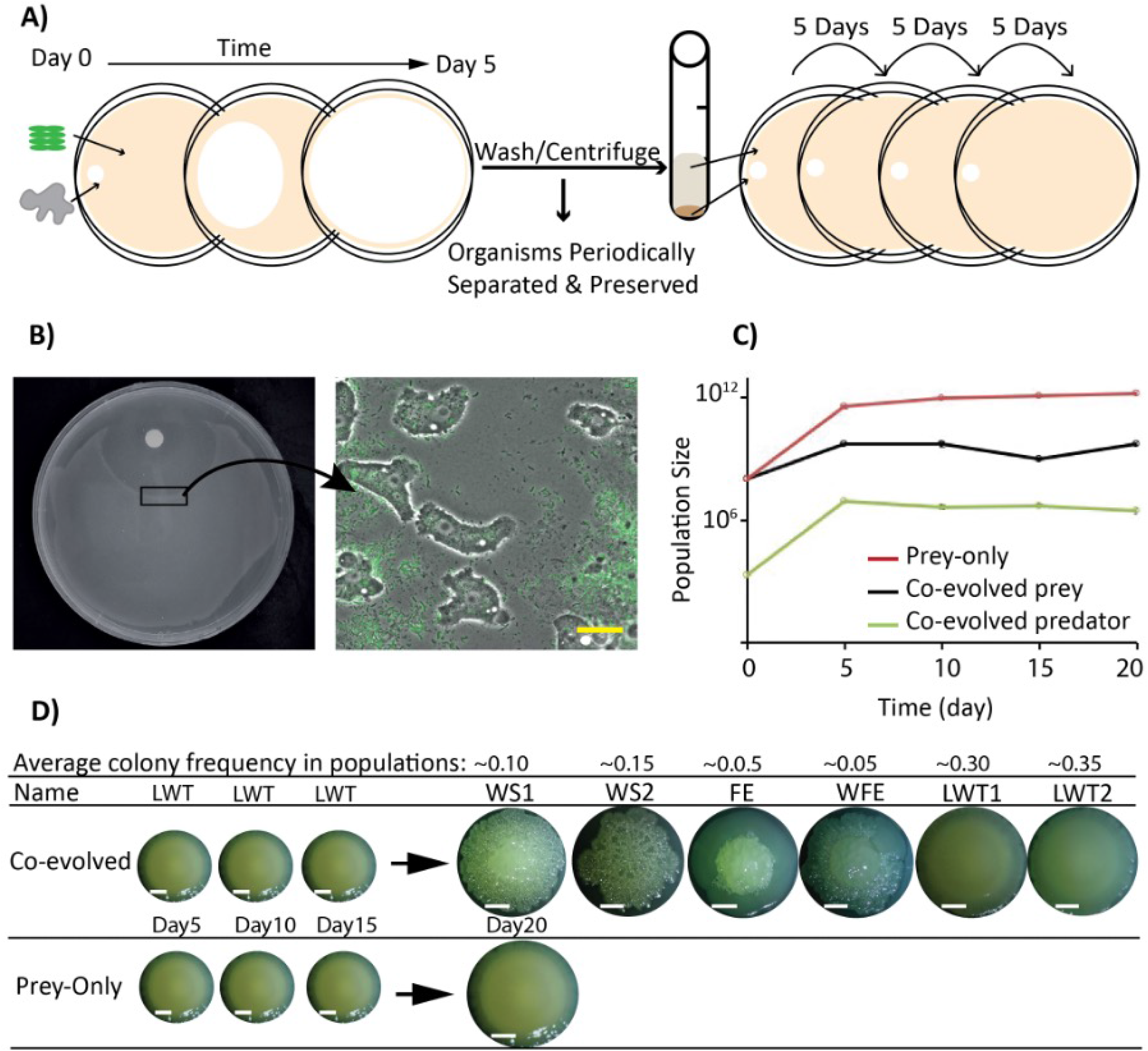
Experimental co-evolution of *P. fluorescens* SBW25 and *N. gruberi* drives bacterial diversification. A) Diagram of the experimental co-evolution protocol on solid media. B) A representative predation-evolution plate. The filter paper, containing ~10^3^ *N. gruberi* can be seen at the top and the progress of the feeding front of amoeba consuming the bacterial lawn can be seen as a decrease in the thickness of the bacterial lawn (left). The feeding front contains active amoeba and the GFP labelled *P. fluorescens* SBW25, visible under fluorescent microscopy (scale bar, 100 μm). C) Estimates of the population size at each transfer for each organism in the predator-associated evolution and the prey-only control over 20-days of evolution. D) Divergence of the bacterial colony morphologies occurs in the presence of predators but not in their absence and the average frequency of the colonies in three evolved populations (scale bar, 10 mm). Colony morphologies pictured are representative of Wrinkly Spreader (WS), Fried Egg (FE), Wrinkly Fried Egg (WFE) and Like Wild Type (LWT). Note, a single representative Smooth, or “like WT” bacterial colony is shown.

**Figure 2.**
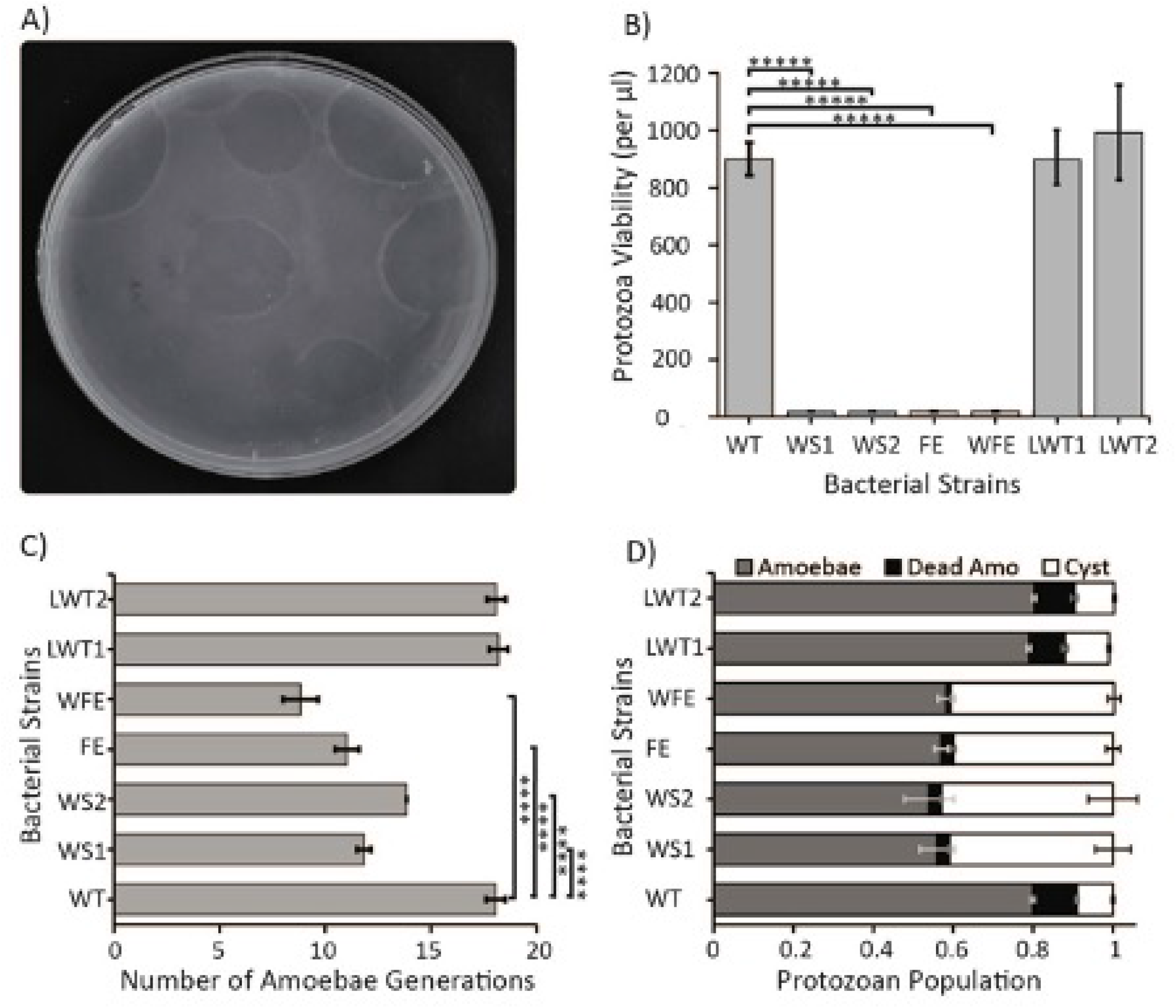
Assays conducted to test the growth and survival of ancestral amoebae obtained on different predator-evolved bacterial isolates. A) *N. gruberi* cells form plaques on a bacterial lawn (each plaque indicative of one amoebae cell) when a single amoeba grows, divides and consumes a large number of bacterial cells over time. B) Estimates of the numbers of *N. gruberi* cells when growing on each bacterial isolate based on plaque counts (n=6, *****P<0.00001, Two sample T tests). C) Average estimation of *N. gruberi* generation time over four days on various evolved bacteria (n=6, ****p≤ 0.00007, Two sample T tests). D) Measuring the approximate number of *N. gruberi* after they were confronted with each of the evolved mutants over four days (n=4). Error bars indicate standard deviation and stars denote statistical significance.

**Figure 3.**
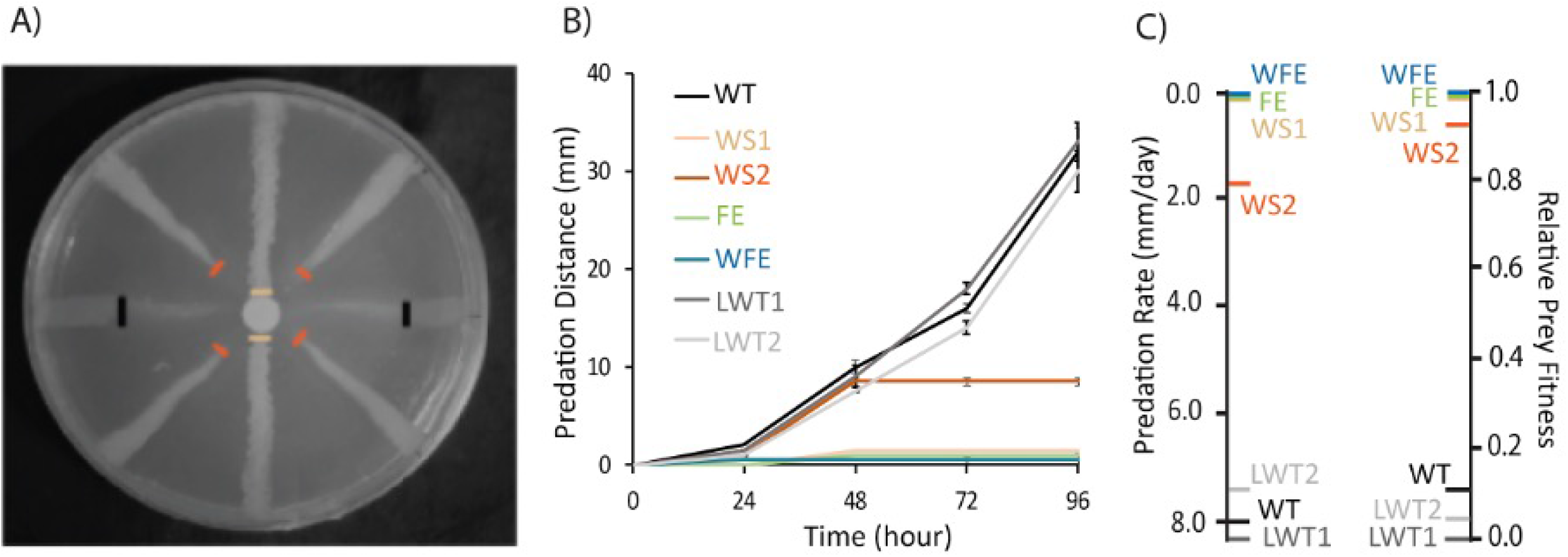
Predation of ancestral *N. gruberi* against evolved *P. fluorescens* SBW25 populations. A) A representative plate showing a line test assay after 72 hours. B) Line test disappearance produced by *N. gruberi* predation. Error bars indicate standard deviation (N=8). C) Rate of *N. gruberi* predation on the colony morphologies of interest as calculated in line tests and relative fitness of prey.

As the evolution under predation progressed, the viable bacterial and amoeboid populations were monitored by plating for CFU and PFU at the time of each transfer (Fig. 1C). The populations of the bacteria and the amoebae remained relatively stable (3 x 10^9^, +/- 2 x 10^9^ and 5.5 x 10^6^, +/- 2.5 x 10^6^, respectively) in all experimental lineages throughout the course of the 20-day experiment and no extinction events were observed.

At each transfer, CFU plates were inspected for colony morphology changes that would be indicative of genetic changes in response to predation pressure. WT colony morphologies were maintained for the first three transfers and it was only upon inspection of the CFU from the fourth transfer that novel morphologies were observed. Colony morphologies on the nopredation plates remained Smooth throughout the experiment. These novel colony morphologies were associated with the appearance of microcolonies in the lawns of *P. fluorescens* SBW25 cells undergoing predation. Four novel colony morphologies and two WT representatives were selected from the predation plates to represent the dominant types observed at the end of the evolution under predation (day 20). These representative colonies were as follows: Wrinkly Spreader 1 and 2 (WS1, WS2), Fried Egg (FE), Wrinkly Fried Egg (WFE) and Like Wild Type 1 and 2 (LWT1, LWT2) (Fig. 1D). These six were isolated, confirmed to be GFP labelled *P. fluorescens* SBW25 cells and preserved at −80°C. Intriguingly, 20-35% of these colony morphologies across the lines appeared to be similar to the “Wrinkly Spreader” colonies observed in the AL adaptive radiation experiments previously described in *P. fluorescens* SBW25 [41].

### Evaluating Growth and Survival of *N. gruberi* on Predation Adapted Bacteria

The evolution of divergent colony morphologies under this predation regime does not guarantee that the bacteria have traits that are either adaptive or associated with increased virulence. We first wanted to establish the degree to which *N. gruberi* were able to grow and subsist on these predation-adapted bacteria. We therefore measured ancestral *N. gruberi* plaque formation on the evolved bacterial isolates of interest and a WT control in a plaque test assay (Fig. 2A).

We did not observe a significant difference in *N. gruberi* plaque formation on the evolved LWT1 and LWT2 isolates compared to the *P. fluorescens* SBW25 WT. However, we did not observe obvious plaque formation when *N. gruberi* were plated on lawns of WS1, WS2, FE or WFE colony morphology mutants (Fig. 2B). Despite the absence of prominent plaques on these isolates, we noted some deformation of the surface that suggested cryptic and possibly reduced predation and growth of the *N. gruberi*. This suggested either that the evolved *P. fluorescens* SBW25 are directly inhibiting robust growth of the predator, or that the *N. gruberi* are unable to efficiently consume the mutant cells in quantities that produce robust observable plaques, or that some plaques were being formed underneath a layer of extracellular material produced by the bacteria.

In order to determine the degree to which predation-evolved isolates limit the strong growth of the *N. gruberi* cells we performed the plaque test assay once again, washing at T=0 and T=4 days in order to estimate the average number of amoebal generations on ancestral or evolved isolates. We found that in the presence of WT or LWT (evolved) bacteria, the *N. gruberi* were able to divide approximately 19 times. However, when *N. gruberi* cells were grown in the same conditions on the evolved *P. fluorescens* SBW25 isolates, they achieved far fewer divisions. *N. gruberi* in the presence of lawns of WS1, WS2, FE, and WFE were only able to divide 11, 14, 10.3 and 9 times, respectively (Fig. 2C). These results suggest that these mutants have a negative influence on replication and division of *N. gruberi* cells leading to the slow growth of protozoan predators.

Reduced division in the presence of evolved prey bacteria could be due to 1) increased cell death, 2) reduced access to bacterial prey as a food source or 3) reduced metabolic activity in an unfavourable environment (increased encystment). In order to evaluate the evidence for these possibilities, we subjected *N. gruberi* to each of the evolved isolates of interest for four days, and measured life cycle stages and viability by flow cytometry (Sup. Fig. 1), using propidium iodide staining to enumerate dead amoeboid cells.

We observed that up to 80% of the *N. gruberi* cells are in the active vegetative form when growing on WT, LWT1 and LWT2 isolates and less than 10% of their population was encysted (Two sample T test, P< 0.02 and P< 0.06). In contrast, more than 40% of the *Naegleria* population was encysted when the prey was WS1, WS2, FE or WFE (Two sample T test, P< 0.002, P< 0.006, P< 0.00006, P< 0.00003) (Fig. 2D). Further, amoebae cells have shown death rates of up to 11%, 9% and 10% in the presence of lawns of WT, LWT1 and LWT2, respectively (Two sample T test, P< 0.01, P< 0.1); 3% death when exposed to the lawns of WS1, WS2 and FE (Two sample T test, P< 0.0001, P< 0.00001, P< 0.0001); and 1% death when growing with the WFE isolate (P< 0.000005) (Fig. 2D).

### Predation on Mutants by *N. gruberi*

The amoebae viability results suggest that an increase in bacterial virulence has not occurred; to the contrary, more dead amoebae are observed in the presence of the ancestral WT and LWT isolates, not the morphologically distinct isolates of interest. The latter increased amoebae encystment rather than death. This suggested to us that our bacteria had adapted in a way that made them unavailable to the protozoan predators. In order to establish the degree to which our bacterial isolates had evolved resistance to grazing by the ancestral *N. gruberi* we examined the speed of predation by performing line-test assays with the evolved isolates and a *P. fluorescens* SBW25 WT control strain as described previously [57] (Fig. 3A). Line tests allow us to measure the predation rate by observing the distance that amoebae have consumed bacteria down a line on the plate (from a filter in the centre). Predation rates on evolved morphotypes LWT1 and LWT2 were very similar to the true WT (Fig. 3B & C). However, the rate of predation by the ancestral *N. gruberi* was reduced when the FE, WS, or WFE mutants were provided. There was a notable difference between WS1 and WS2: the latter was consumed early in the line test assay and then the *N. gruberi* appeared to have stopped consuming these lines (Fig. 3B & C).

### Bacterial Fitness on Solid Surfaces

In order to understand the fitness cost of the mutations in evolved bacterial isolates we estimated bacterial growth in the presence and absence of the protozoan predator on agar plates. WT and LWTs reached higher CFU over four days than the evolved isolates. As expected, mutants that confer resistance (WS1, WS2, FE and WFE) are more able than WT to withstand predation and experience more generations in the presence of predators (Fig. 4A & B). However, in the absence of predators the ability of three of these mutants (WS1, WS2 and WFE) to replicate and grow were reduced (Fig. 4A) and their average generations on plates over 4 days were similarly decreased (Fig. 4B). This suggests that evolved predation resistance imposes a fitness cost in this environment. Interestingly, we did not measure a significant trade-off in the FE mutants.

**Figure 4.**
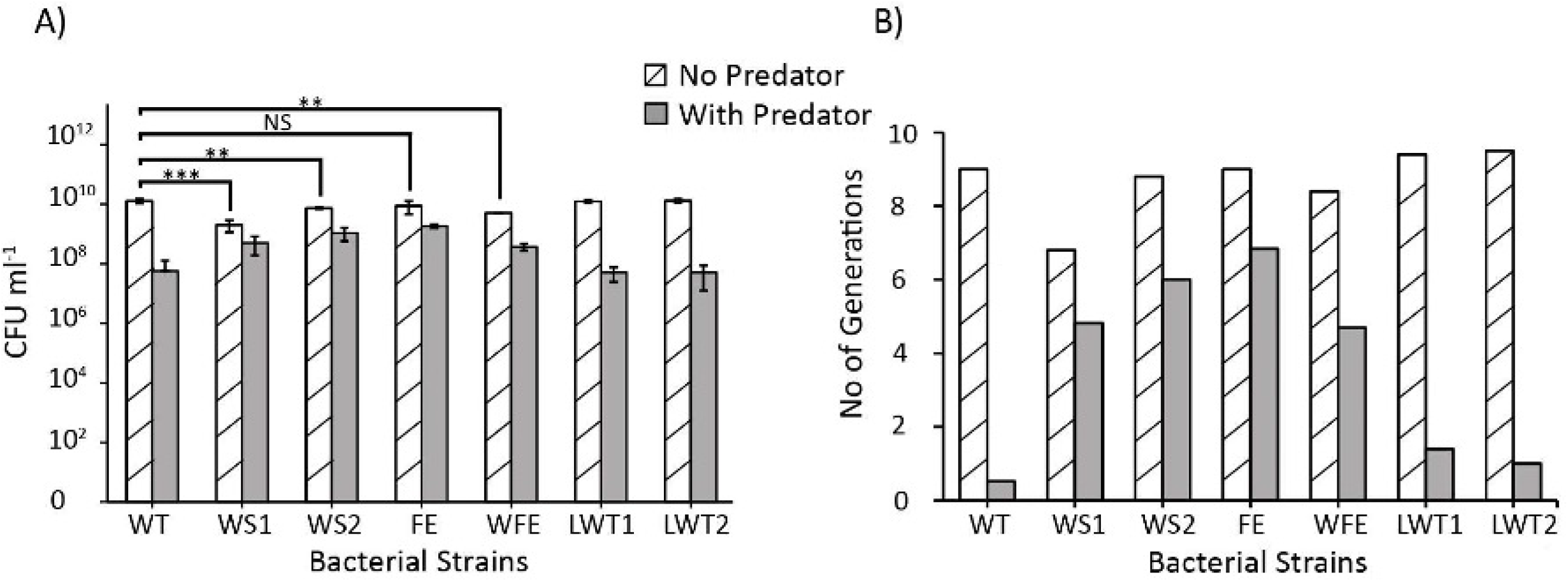
Bacterial fitness on agar surface with and without ancestral predators. Bacteria were grown in the presence and absence of predator *N. gruberi* for four days. (N=4, ***P<=0.002, **P<0.02, NS = not significant). Error bars indicate standard deviation and stars denote statistical significance.

### Sequencing Results

The colony morphologies that we observed after 20 days of evolution under protozoan predation are reminiscent of the WS mutations observed at the AL interface in *P. fluorescens* SBW25. In order to determine how similar the underlying mutations might be, we subjected the evolved isolates and three WT colonies from the non-predation plates to shotgun DNA sequencing.

Five of the six evolved isolates each had a single unique mutation (Table 1). The exception is the LWT2 isolate, for which no mutations were detected. Moreover, three of the six evolved isolates had acquired mutations in *wspF* (PFLU_1224). Mutations in *wspF* are frequently observed in studies of *P. fluorescens* SBW25 during adaptive radiation at AL interfaces and are consistent with the “wrinkly” colony morphology observed (Fig. 5A, Sup. table 1) [61].

**Table 1.** The locations and details of mutations observed in this study by whole-genome sequencing.

| Isolate | Gene name | PFLU | ORF Nucleotide<br>Change | ORF AA<br>Change | Domain | Morph |
| --- | --- | --- | --- | --- | --- | --- |
| SBW25 | — | — | — | — | — | Smooth |
| WS1 | *wspF* | 1224 | Δ231-236 | ΔVIV 76-78 | CheY<br>phosphorylation | Wrinkly<br>Spreader |
| WS2 | *wspF* | 1224 | +166-180 | +LMDLI 56-60 | CheY<br>phosphorylation | Wrinkly<br>Spreader |
| FE | *wspF* | 1224 | T815C | L272P | CheB-type<br>methylesterase | Fried Egg |
| WFE | *amrZ/algZ* | 4744 | C97G | A33P | DNA<br>binding | Wrinkly Fried<br>Egg |
| LWT1 | *gcvA; LysR* | 3329-30 | ΔTGGGCCACC | — | NA | Like Wild Type |
| LWT2 | No mutation | — | — | — | — | Like Wild Type |

**Figure 5.**
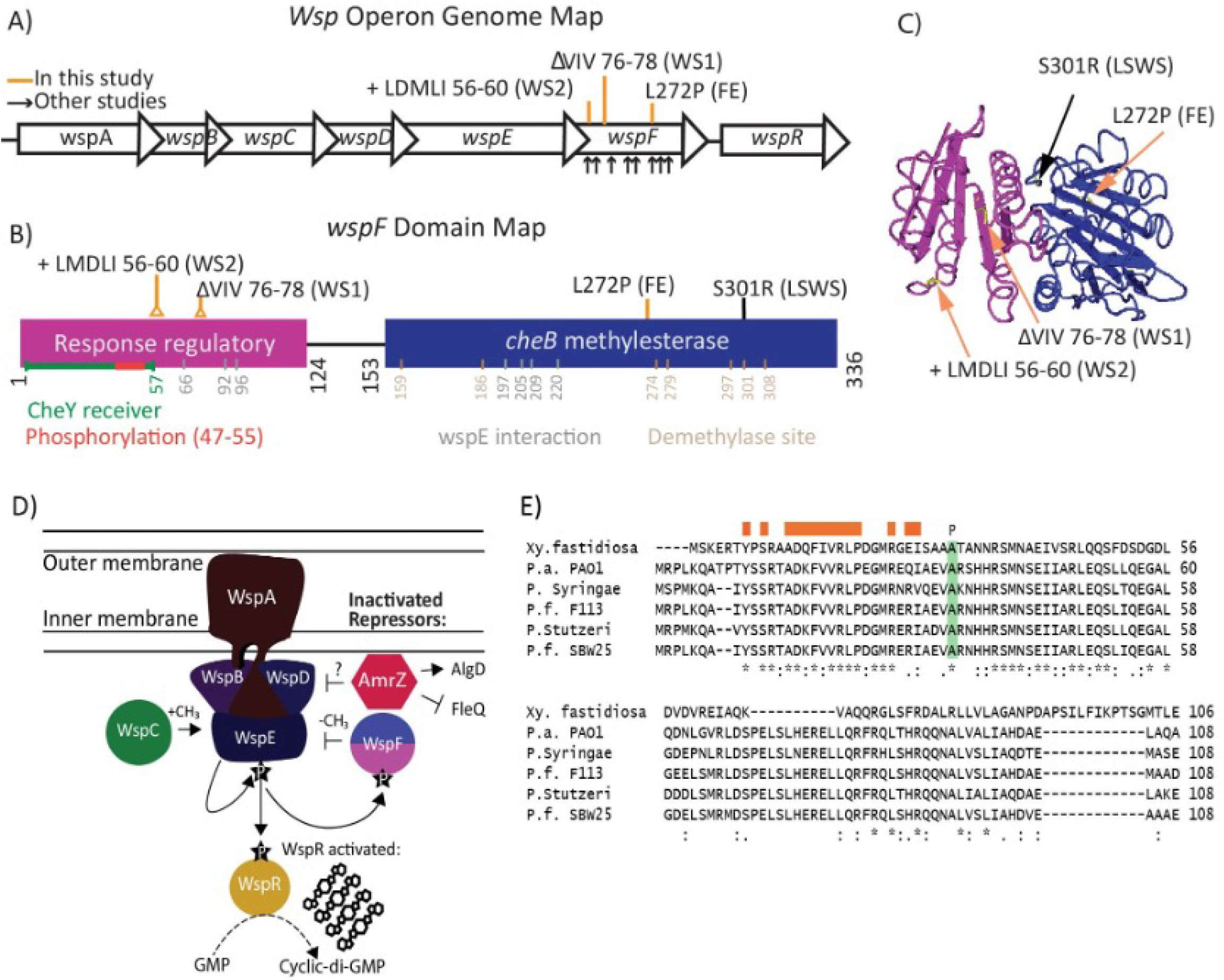
Amoeboid predation resistance can be conferred by biofilm formation in *P. fluorescens* SBW25. A) The *wsp* operon in *P. fluorescens* SBW25, the *wspF* mutations found in this study (orange lines) resulting in Wrinkly Spreader phenotypes and those from AL interface studies (black arrows). B) The *wspF* domain map in *P. fluorescens* SBW25; predicted sites and the mutations identified in this study are shown above the domain map plus the LSWS positive control. C) WspF protein structure in SBW25 adopted from *Salmonella Typhimurium* (PDP ID; 1A2O). The protein consists of two domains: the CheB-type methylesterase domain (blue) and a response regulator-like *cheY* receiver domain in pink. Predicted mutational targets from this study and LSWS are highlighted in yellow and their names are shown (orange arrows). D) A model describing mutations that conferred resistance to amoeboid predation in *P. fluorescens* SBW25 during a short term co-evolution experiment. FE mutation is in the methyltransferase domain and the WS mutations are both in the CheY domain. E) Amino acid alignment of AmrZ homologs in *P. fluorescens* SBW25 and other bacterial species. The orange box shows the dimerization domain of AmrZ and a mutation resulting in the WFE phenotype is shown in green.

The mutations in Isolates WS1 and WS2 were in the CheY receiver domain of WspF (Fig. 5) [43]. The WS2 mutation was immediately adjacent to the phosphorylation domain within this receiver domain. The mutation in the WS2 isolate overlapped with previously identified *wspF* mutations that had arisen as constitutive cellulose producers in the AL interface experiments due to interruption of the receiver domain of WspF (Fig. 5 and Sup. table 1) [62,63].

The FE isolate was also found to have acquired a mutation in the *wspF* gene, a single nucleotide change affecting the methyltransferase region of the CheB-type methylesterase domain (Table 1, Fig. 5). This mutation is within 2 AA of a previously detected mutation at 274 discovered in the AL interface experiment (Sup. table 1) to lead to a robust WS morphology and constitutive expression of cellulose [43].

The WFE mutant isolate included a single nucleotide change in the *amrZ* ORF (alginate and motility regulator). The AmrZ DNA binding protein has been implicated as a negative regulator of the Wsp operon [52,64], a positive regulator of algD (alginate synthesis) and a negative regulator of flagellum synthesis in *P. fluorescens* (Table 1, Fig. 5D) [64]. To determine the degree to which the *amrZ* mutation might be disrupting function, we aligned the AA sequence of *amrZ* from *P. fluorescens* SBW25 and five other bacterial species [65] (Fig. 5E). The AA identity at position 33 in *amrZ* is a highly conserved residue that lies outside of the dimerization zone. This alignment suggests that the mutation will cause some loss of function.

Capsule formation is a common virulence trait that has been implicated in parasite resistance in *P. fluorescens* SBW25 [66]. We therefore tested all isolates for capsule production by India ink staining and all were negative (Sup. Fig. 2). The LWT1 isolate, despite having acquired a 12-bp deletion in a regulatory region between LysE family translocator *lysR* and transcriptional regulator *gcvA*, showed no change in either colony morphology or predation resistance; thus the deletion is probably neutral in this context (Table 1).

### Swarming test

AmrZ is known to be a negative regulator for flagellum biosynthesis in some *Pseudomonads* [65]. To determine whether the AmrZ mutation affects motility we carried out a swarming test assay, comparing it to WT and ΔFleQ as a negative control [67]. The WFE mutant AmrZ (A33P) showed a swarming defect compared to the WT *P. fluorescens* SBW25 that appeared to be mediated by obvious biofilm formation on the surface of the swarming plate (Fig. 6). The WFE mutant did not demonstrate a greater swarming distance than the △FleQ negative control in the first 21 hours. However, the swarming ability of the WFE mutant appeared to improve after the initial biofilm formation stage, when more cells appeared to breach the surface.

**Figure 6.**
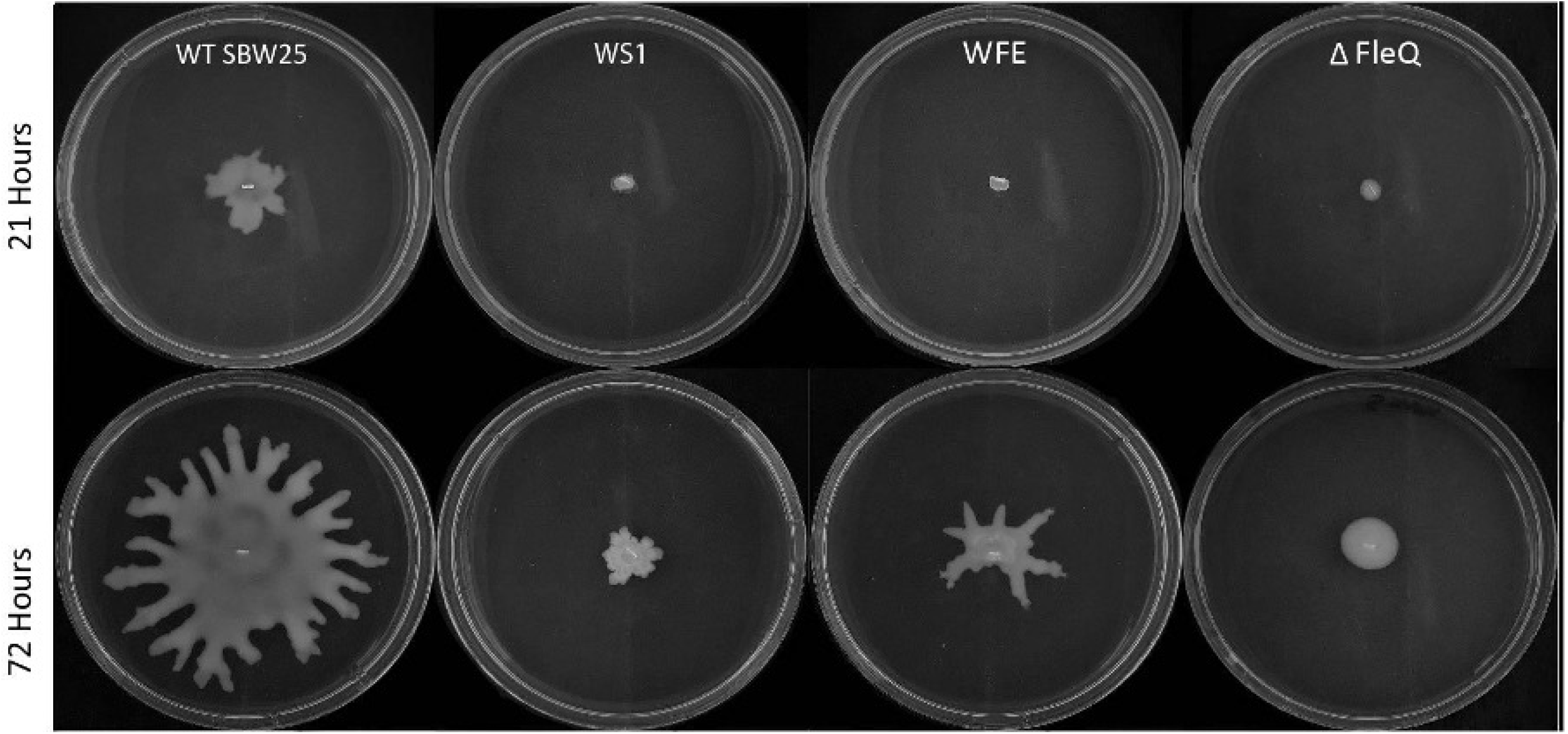
Representative swarming images of the WT, WS1, WFE and FleQ mutants after 21 and 72 hours of growth on M9 plates.

### Mat Strength & Cellulose Expression

Three out of the four colony morphology mutants that we chose to focus on are mutations that are similar but not identical to those observed in the classic AL interface experiments [44]. We therefore examined cellulose production on solid surfaces (Fig. 7A, B and C) and adopted the glass bead strength test to measure the strength of biofilms formed (Fig. 6D) [61]. We included the *P. fluorescens* SBW25 Large Spreading Wrinkly Spreader (LSWS) mutant as a positive control, as this mutant forms rapid and robust biofilms at the AL interface [43]. We initiated KB AL interfaces in standard microcosms from the colony morphology mutants of interest (shown) [44].

**Figure 7.**
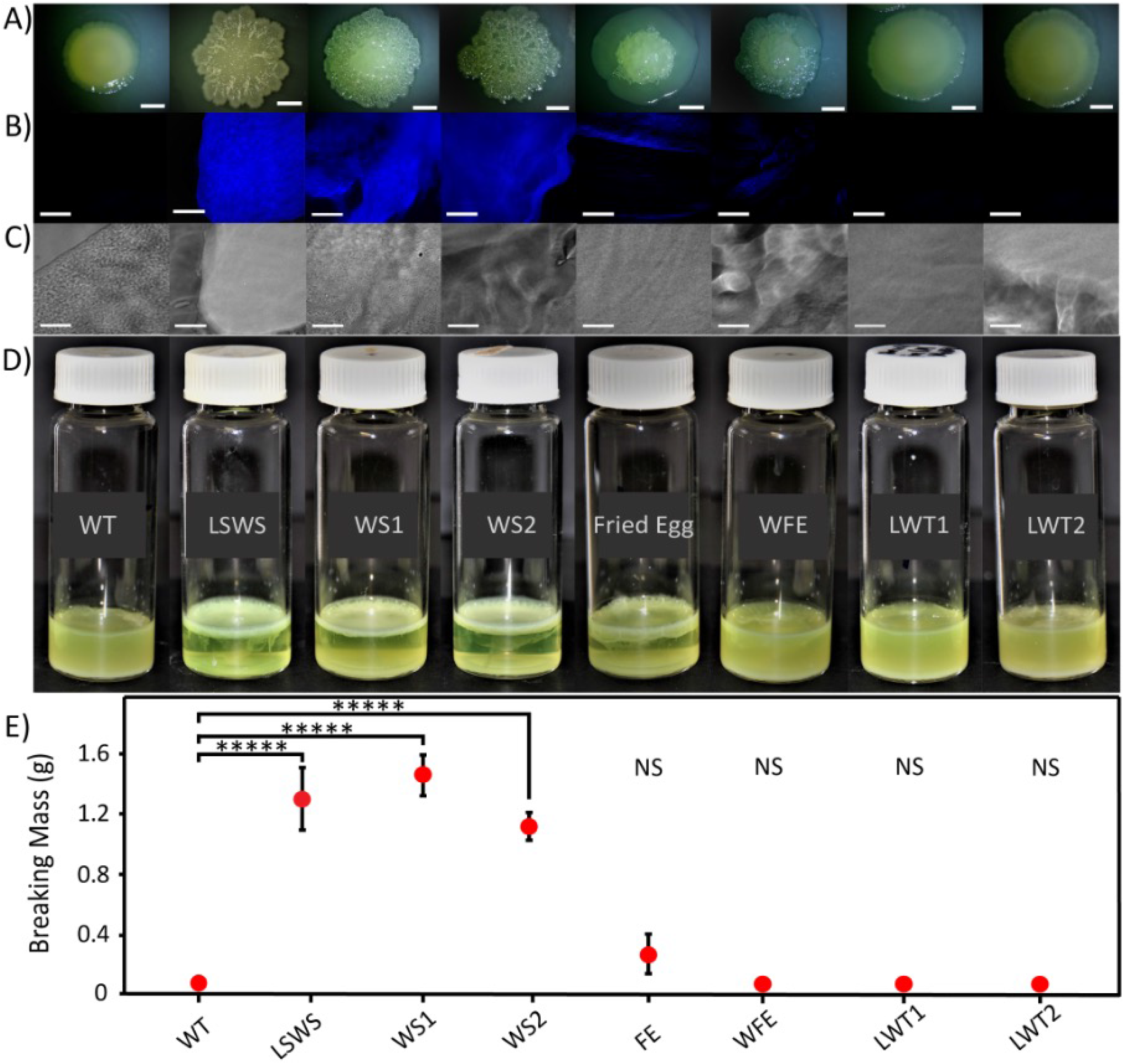
Bacteria evolved under predation develop biofilm morphologies that are comparable to adaptive divergence phenotypes. A) Colony morphologies on LBA (Scale size: 10 mm). B) Visualisation of cellulose production grown on KB agar with a combination of calcofluor staining and microscopy. Scale bar: 30μm. C) Phase contrast image of colony segments shown in figure B. D) Microcosms after three days of growth in KB. E) Graph showing the summed mass of the beads that were added before the biofilm mat broke for each mutant type of interest (N=4, *****P<=0.0005, P values of over 0.05 were deemed insignificant (NS). All P values were pairwise T tests to WT. Error bars indicate standard deviation and stars denote statistical significance.

Within 24 hours, the positive control and four of our mutants of interest occupied the AL interface. As expected, LWT1, LWT2 and *P. fluorescens* SBW25 WT did not readily form biofilms over this time period. The strength of mats formed by WS1 and WS2 (1.3 g +/- 0.18) were similar in strength to the LSWS (1.1 g +/- 0.4). These mutants produced significantly stronger mats than the WT (Fig. 7D & E). The FE mutant and WFE mutants were not as successful at colonizing the AL interface and the mats produced were significantly less robust (0.15 g +/- 0.15), suggesting that these morphotypes are not robust biofilm producers. Calcofluor white staining confirmed the AL interface findings [68]. WS1 and WS2 each produced abundant extracellular cellulose whilst FE and WFE showed much less extracellular cellulose production (Fig. 7B).

## Discussion

We conducted a 20-day predator-prey co-evolution experiment in order to experimentally evaluate bacterial adaptations that increase resistance to amoeboid predation on solid surfaces. The predation-free lines did not develop recognizable colony variants, but after 20 days of predation a set of novel and not-so-novel bacterial colony morphologies were selected from the predation adapted bacterial lineages for further study. Two of the novel types closely resembled WS types observed in AL experiments performed previously in the same organism while the other two, WFE and FE, were not previously described.

All four colony variants demonstrated increased grazing resistance in plaque assays and line test assays and all four had substantially increased prey fitness relative to the WT. The strategies in place to resist predation appear however to be distinguishable.

As noted, two of the colony variants selected were classified as WS types. Complete genome sequencing revealed that the WS1 and WS2 isolates both had mutations in the phosphorylation domain of the well characterized *wspF* gene (Fig. 5). Similar mutations in this response regulatory domain are well known to increase cellular levels of cyclic-di-GMP and ultimately cause cells to constitutively over-produce extracellular cellulose, resulting in robust biofilms at the AL interface. In light of this genotype we tested WS1 and WS2 in a classic AL interface experiment and found they produced high strength biofilms, comparable to a well studied positive control (LSWS) [52]. This extracellular production strategy parallels the bacterial prey *Escherichia coli* and bacterial predator *Myxococcus xanthus* predation experiments that favour increased extracellular mucoid production [69]. The evolution of constitutive biofilm production as an adaptive response to predation is consistent with expectations [18]. There is growing evidence that biofilm formation is a common response of *Pseudomonad*s to ecological challenges, aside from protozoan predation, including AL interfaces [44], the CF lung [70], and even space travel [71]. This may be an ecological consequence of the many negative repressors that contribute to the regulation of this trait, providing a large target for random mutations [52].

We observed trade-offs in the fitness of the majority of bacterial mutants that had acquired resistance to amoeboid predation (Fig. 4B). This was particularly pronounced in the WS1 mutant, which lost an average of 23.8% generational growth compared to WT on these plates. The relative fitness of WS morphotypes measured in mixed colony fitness assays was 0.33 of WT over 100 generations [72]. In our hands, WT cells achieved an average of 9 generations over the 4-day experiment. A loss of 23.8% over a short time represents a substantial trade-off. However, trade-offs of this kind were not universal in these predation resistant mutants.

The FE mutation did not appear to impose a selective cost as measured by cell growth on solid media in the absence of predators (Fig. 4B). This may suggest that the strategy of these cells to resist predation is optimal. It is also possible that we have failed to measure their growth in an environment in which deleterious fitness effects of this *wspF* mutation would be apparent.

The FE colony variant was also revealed to be a *wspF* mutation, however this one was in the demethylase domain of the WspF repressor. This mutation (L272P) is comparable to the LSWS mutation, which is found in the same domain (Fig. 5B & C). The result is a much less robust biofilm, which can only maintain the weight of 0.31 grams whereas the LSWS can support 1.33 grams. The cellulose production was also much less pronounced in the FE isolate as visualized by calcofluor staining (Fig. 7). This mutation is proximal (2 AA away) from the WSF (AA274) mutation observed previously in AL experiments (Sup. Table 1) and shown to have a higher fitness than the LSWS in AL competitions [62]. Our results suggest that the FE mutation modulates the methylase activity of the WspF domain but the effect on the balance of Wsp and therefore c-di-GMP accumulation in the cell and cellulose production are not as dramatic as in the LSWS (S301R) mutation in the same domain [62].

The WFE colony variant also affects the function of a repressor that is implicated in c-di-GMP production via the Wsp regulatory system but this mutation was found in the *amrZ* gene, a transcriptional regulator. AmrZ is known to repress the Wsp system whilst simultaneously stimulating the production of alginate through AlgD and repressing production of the flagella through FleQ (Fig. 5D & E) [52,64]. In our hands, the mutation in AmrZ appeared to have a mild effect (similar to that of FE) in stimulating cellulose production, manifest as poor performance in the AL interface and limited calcofluor staining (Fig. 7). We interpret this as a partial loss of function of the AmrZ repressor. We have no evidence that this mutation increases flagella production concomitant with the observed increase in cellulose production [64]. Our swarming motility assay result suggests that motility is impaired but this may be a consequence of cellulose production (Fig. 6). These somewhat contradictory phenotypes, increased motility and increased biofilm formation, may explain the extremely weak biofilm formed in this instance.

We set out to determine if evolution under amoeboid predation would lead to an increase in bacterial virulence, focussing on altered colony morphologies in order to identify mutations. It is perhaps unsurprising that the isolates characterised were those involved in biofilm formation to one degree or another. While biofilm formation has certainly been implicated as a virulence trait in some *Pseudomonads* [14,70], the WS1, WS2, FE and WFE variants were highly resistant to predation without increasing amoeboid mortality.

Contrary to our expectations, we found that the *N. gruberi* that preyed on LWT or WT isolates had higher mortality (10% cell death) than those consuming the WS, FE or WFE variants (Fig. 2D). *N. gruberi* demonstrated significantly better growth (19 generations) in the presence of the WT-like isolates than they did in colony variants (9-14 generations). This increased mortality of growing *N. gruberi* likely results from the defensive traits expressed by WT *P. fluorescens* SBW25 in the presence of *Naegleria*. For example, WT *P. fluorescens* SBW25 has been shown to produce viscosin, a cyclic lipopeptide biosurfactant implicated in increased motility and surface spreading and which has lethal effects on *N. americana* [67,73]. We interpret the decreased mortality observed in the presence of bacterial biofilms to suggest this presents a food deprivation response in which *N. gruberi* revert to their encysted state, reducing their exposure to bacterial defenses and thereby reducing mortality [56]. Given the antagonistic starting point for our predator and prey, longer co-evolution experiments may yield isolates that increase in virulence, particularly given the tendency of experimental evolution protocols that require subculturing and transfer to select against biofilm-forming mutants in the long term [7].

Our findings support the suggestion that amoeboid predation can profoundly influence the course of genetic and phenotypic evolution in a short time span. Protozoan predation is an important, but often neglected, determinant of bacterial populations in nature [6]. The power of these bacterial antagonists has been recognized previously but there has been a dearth of studies on the long-term genetic effects these keystone predators have on bacterial prey. FLAs can be integral modulators of the bacterial community in biofilms [74,75]. FLAs isolated from several environmental samples were recognised often to be in close contact with the bacterial biofilms in nature [4]. Biofilm formation has previously been demonstrated to be a successful defence strategy of *P. fluorescens* SBW25 against free-living ciliate predators in liquid environments [76–78]. Future work with this model system may be able to support or refute the hypothesis that protozoan predation has the potential to drive the evolution of virulence traits in bacteria.

## Supporting information

Supplemental File All

## Acknowledgements

We are grateful to Xue-Xian Zhang for kindly providing the *P. fluorescens* SBW25 *ΔfleQ* mutant.

## Author Contributions

HH designed the experiments in collaboration with FG. FG contributed to the writing of the first draft of the manuscript. GF and HH contributed to manuscript reading and revision.

---

**Supplemental Figure 1**. Results of flow cytometry showing *N. gruberi* cell populations distinguished by morphology and viability using PE-A and FSC-A channels. Active amoeba, cyst and dead populations are shown in green, blue and red, respectively.

**Supplemental Figure 2**. Capsule Staining We examined the predator-evolved phenotypes for production of a colanic acid-like polymer [79] by visualising them with India Ink (see materials and methods) and observed them using phase contrast microscopy. We included a positive control named 6B4 [45] that makes large capsules. None of the mutants were able to produce a capsule. There were no mutations found in any of our predation-adapted genotypes in known encapsulation associated genes and the evolved isolates do not appear to have increased capsule formation under standard laboratory conditions.

