## Supplemental File All for "Protozoan predation drives adaptive divergence in *Pseudomonas fluorescens* SBW25; ecology meets experimental evolution"

**Supplemental tables and figures for Protozoan predation drives adaptive divergence in *Pseudomonas fluorescens* SBW25; ecology meets experimental evolution.**

Farhad S. Golzar^1^, Gayle C. Ferguson^1^, and Heather Lyn Hendrickson^1^

**Supplemental Table 1** *wspF* mutations associated with Wrinkly Spreader Phenotype in this study and the previous literature.

| **WS genotype** | **Gene** | **Nucleotide change** | **AA change** | **Reference** |
| --- | --- | --- | --- | --- |
| WS1 | *wspF* | ∆231-236 | ∆VIV 76-78V | This study |
| WS2 | *wspF* | +166-180 | +LMDLI 56-60 | This study |
| FE | *wspF* | T815C | L272P | This study |
| LSWS | *wspF* | A901C | S301R | [[62]](https://paperpile.com/c/w3Y1c7/VbPt) |
| WSA | *wspF* | T14G | I5S | [[62]](https://paperpile.com/c/w3Y1c7/VbPt) |
| WSB | *wspF* | ∆620-674 | P206∆ | [[62]](https://paperpile.com/c/w3Y1c7/VbPt) |
| WSC | *wspF* | G823T | G275C | [[62]](https://paperpile.com/c/w3Y1c7/VbPt) |
| WSE | *wspF* | G658T | V220L | [[62]](https://paperpile.com/c/w3Y1c7/VbPt) |
| WSF | *wspF* | C821T | T274I | [[62]](https://paperpile.com/c/w3Y1c7/VbPt) |
| WSG | *wspF* | C556T | H186Y | [[62]](https://paperpile.com/c/w3Y1c7/VbPt) |
| WSJ | *wspF* | ∆865-868 | R288∆ | [[62]](https://paperpile.com/c/w3Y1c7/VbPt) |
| WSL | *wspF* | G482A | G161∆ | [[62]](https://paperpile.com/c/w3Y1c7/VbPt) |
| WSN | *wspF* | A901C | S301R | [[62]](https://paperpile.com/c/w3Y1c7/VbPt) |
| WSO | *wspF* | ∆235-249 | V79∆ | [[62]](https://paperpile.com/c/w3Y1c7/VbPt) |
| WSU | *wspF* | ∆823-824 | T274∆ | [[62]](https://paperpile.com/c/w3Y1c7/VbPt) |
| WSW | *wspF* | ∆149 | L49∆ | [[62]](https://paperpile.com/c/w3Y1c7/VbPt) |
| WSY | *wspF* | ∆166-180 | ∆L51-I55 | [[62]](https://paperpile.com/c/w3Y1c7/VbPt) |
| WS | *wspF* | - | 0297K | [[80]](https://paperpile.com/c/w3Y1c7/jLcT) |
| WS | *wspF* | - | V271G | [[80]](https://paperpile.com/c/w3Y1c7/jLcT) |
| WS | *wspF* | - | G270R | [[80]](https://paperpile.com/c/w3Y1c7/jLcT) |
| WS | *wspF* | - | P47L | [[81]](https://paperpile.com/c/w3Y1c7/9ncj) |
| WS | *wspF* | - | ∆R66-L 107 | [[81]](https://paperpile.com/c/w3Y1c7/9ncj) |
| WS | *wspF* | - | S159L | [[81]](https://paperpile.com/c/w3Y1c7/9ncj) |
| WS | *wspF* | - | H186Y | [[81]](https://paperpile.com/c/w3Y1c7/9ncj) |
| WS | *wspF* | - | Q297R | [[81]](https://paperpile.com/c/w3Y1c7/9ncj) |
| WS | *wspF* | - | ∆T226-G275 | [[81]](https://paperpile.com/c/w3Y1c7/9ncj) |

Supplemental Figures

Supp-Fig-1

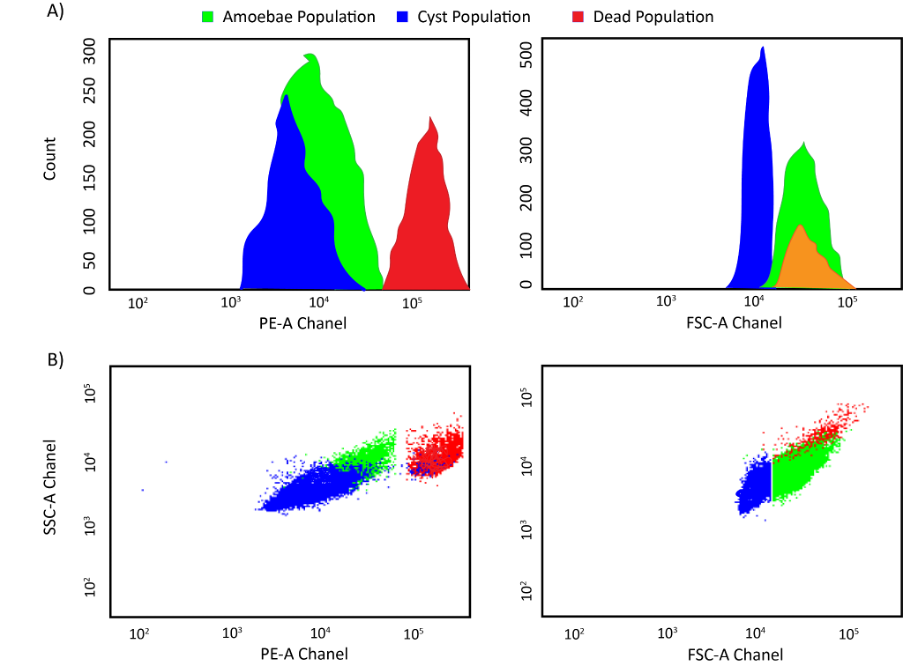

**Supplemental Figure 1**. Results of flow cytometry showing *N. gruberi* cell populations distinguished by morphology and viability using PE-A and FSC-A channels. Active amoeba, cyst and dead populations are shown in green, blue and red, respectively.

Supp-Fig-2

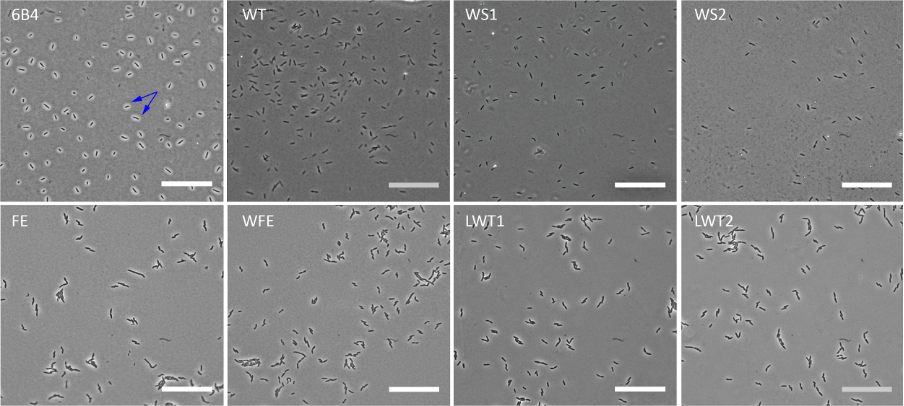

**Supplemental Figure 2**. Capsule Staining

We examined the predator-evolved phenotypes for production of a colanic acid-like polymer [[79]](https://paperpile.com/c/w3Y1c7/dDd5) by visualising them with India Ink (see materials and methods) and observed them using phase contrast microscopy**.** We included a positive control named 6B4 [[45]](https://paperpile.com/c/w3Y1c7/ekNq) that makes large capsules. None of the mutants were able to produce a capsule. There were no mutations found in any of our predation-adapted genotypes in known encapsulation associated genes and the evolved isolates do not appear to have increased capsule formation under standard laboratory conditions.
